# Engineering targeted deletions in the mitochondrial genome

**DOI:** 10.1101/287342

**Authors:** Jarryd M. Campbell, Ester Perales-Clemente, Hirotaka Ata, Noemi Vidal-Folch, Weibin Liu, Karl J. Clark, Xiaolei Xu, Devin Oglesbee, Timothy J. Nelson, Stephen C. Ekker

## Abstract

Mitochondria are a network of critical intracellular organelles with diverse functions ranging from energy production to cell signaling. The mitochondrial genome (mtDNA) consists of 37 genes that support oxidative phosphorylation and are prone to dysfunction that can lead to currently untreatable diseases. Further characterization of mtDNA gene function and creation of more accurate models of human disease will require the ability to engineer precise genomic sequence modifications. To date, mtDNA has been inaccessible to direct modification using traditional genome engineering tools due to unique DNA repair contexts in mitochondria^1^. Here, we report a new DNA modification process using sequence-specific transcription activator-like effector (TALE) proteins to manipulate mtDNA *in vivo* and *in vitro* for reverse genetics applications. First, we show mtDNA deletions can be induced in *Danio rerio* (zebrafish) using site-directed mitoTALE-nickases (mito-nickases). Using this approach, the protein-encoding mtDNA gene *nd4* was deleted in injected zebrafish embryos. Furthermore, this DNA engineering system recreated a large deletion spanning from *nd5* to *atp8*, which is commonly found in human diseases like Kearns-Sayre syndrome (KSS) and Pearson syndrome. Enrichment of mtDNA-deleted genomes was achieved using targeted mitoTALE-nucleases (mitoTALENs) by co-delivering both mito-nickases and mitoTALENs into zebrafish embryos. This combined approach yielded deletions in over 90% of injected animals, which were maintained through adulthood in various tissues. Subsequently, we confirmed that large, targeted deletions could be induced with this approach in human cells. In addition, we show that, when provided with a single nick on the mtDNA light strand, the binding of a terminal TALE protein alone at the intended recombination site is sufficient for deletion induction. This “block and nick” approach yielded engineered mitochondrial molecules with single nucleotide precision using two different targeted deletion sites. This precise seeding method to engineer mtDNA variants is a critical step for the exploration of mtDNA function and for creating new cellular and animal models of mitochondrial disease.

## Text

Mitochondria play a pivotal role in many cellular processes; including, ATP production, ion homeostasis, iron-sulfur cluster biogenesis, cell signaling and, when deficient, cause divergent and progressive conditions^2,3^. Severe defects in mtDNA result in debilitating, incurable disorders, and manipulation of mtDNA for disease modeling has been largely limited to either random mutagenesis^4–6^ or naturally occurring alterations. Moreover, severe mtDNA variants do not readily pass through the germline^7^, making it difficult to develop stable animal models of more severe, physiologically-relevant diseases. Indeed, understanding of this critical organelle in health and disease has been stymied by our inability to precisely manipulate mtDNA *de novo*. Standard editing tools like TALENs and CRISPR-Cas9 primarily rely on double-strand break (DSB) repair in the nuclear genome, where both non-homologous end joining (NHEJ) and homology directed repair (HDR) are active. Importantly, DSBs are known to trigger rapid mtDNA degradation and have been used strategically to combat the deleterious effects of heteroplasmy in patient cells and mouse models of mtDNA disease^8–12^. Although DSBs can be repaired at low frequencies in mitochondria^13–15^, both NHEJ and HDR are very inefficient and not useful to date for mtDNA sequence engineering^16^.

To address this technical challenge, we developed a novel system for targeted manipulation of mtDNA sequences using alternative editing strategies. One option for mtDNA targeting includes CRISPR/Cas9; however, targeting to mitochondria would require effective gRNA importation, a hurdle yet unresolved^1,17^. TALE proteins are traditionally more difficult to assemble than CRISPR/Cas9, but single-reaction synthesis technology makes TALEs readily accessible for genome engineering applications^18^. Importantly, TALE use in mtDNA engineering only require a single mitochondrial targeting sequence (S Figure 1) that was developed from an *in vivo* protein trapping method^19^. We reasoned a nickase-induced single-strand break approach could generate targeted deletions of various sizes in mtDNA by avoiding DSBs^20^. We deployed zebrafish as a rapid *in vivo* initial test system because of its conserved mtDNA genome^21^ and ease of microinjection for efficient and quantitative delivery of genome editing tools (Figure 1A). The zebrafish mitochondrial targeting sequence (S Figure 1) was fused to the GoldyTALEN backbone^22^ in both nuclease and nickase forms, the latter made by mutagenizing half of the FokI dimer rendering it catalytically inactive (p.D450A^23^, Figure 1B). We hypothesized that the resulting mito-nickase could be used to directly manipulate mtDNA genomes for new single-gene deletions as well as site-specific, multigene variants that can molecularly model human disease.

**Figure 1.**
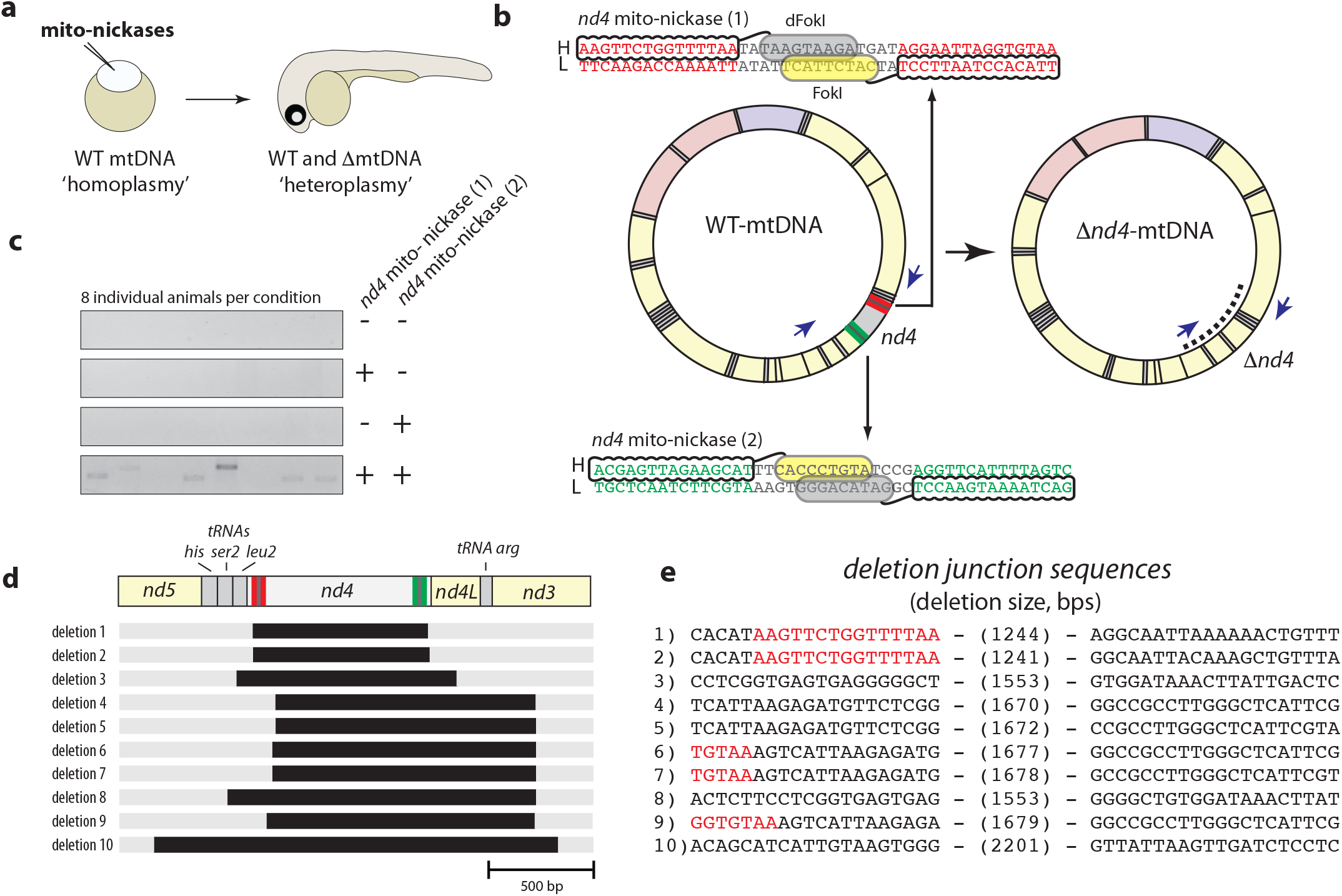
Single-gene deletion of *nd4* using mito-nickases. **a** Schematic of mtDNA sequence manipulation using mito-nickases in zebrafish. **b** Mito-nickases were targeted inside the border of *nd4*, with TALE binding and FokI activity on either the heavy (H) or light (L) strand. The active FokI is in yellow while the catalytically inactive variant, dead FokI (dFokI), is in grey. Deletions were detected using PCR, with primers positioned outside of both nick sites (blue arrows). Expected PCR product indicated by dashed line. **c** Deletion of *nd4* was detected using PCR in animals injected with *nd4* mito-nickases. Every row contains 8 individual animals per condition. **d** Schematic of mtDNA region surrounding *nd4* and corresponding location of deletions shown as black bars. Position of *nd4* mito-nickase (1) is in red, and position of *nd4* mito-nickase (2) is in green. **e** Deletion junction sequences with the size of deletion in parentheses. Red sequences are the *nd4* mito-nickase (1) binding sites.

The 1.2 kilobase, protein-encoding mtDNA gene, *nd4*, was used to test whether our approach could yield deletions for mitochondrial DNA functional testing (Figure 1B). By positioning two mito-nickases just inside the *nd4* coding region, deletions were readily detected in about 50% of injected embryos (Figure 1C). We subsequently sequenced and confirmed that mtDNA deletions were successfully targeted to, or near, the mito-nickase sites (Figure 1D,E). In every deletion screened, *nd4* was deleted along with up to 500 basepairs of adjacent nickase site-sequence (Figure 1D).

Numerous human disorders are caused by mtDNA deletions^2^. For instance, a ‘common deletion’ of 4977 basepairs between *nd5* and *atp8* is identified in most cases of Kearns-Sayre Syndrome (KSS)^24,25^. To molecularly model this deletion, two mito-nickases were targeted within these genes and resulting products were detected by bridge-PCR and sequencing (Figure 2A). The bridge-PCR was designed to selectively amplify mtDNA deletions by positioning two primers outside mito-nickase sites and maintaining a short extension time to avoid amplification of the full-length wild type product (Figure 2A). Using this method, deletions were detected in nearly 30% of injected animals, and sequence verification confirmed deletion junctions near the mito-nickase targeted sites (designated the Δ*nd5*/*atp8*-mtDNA genome, Figure 2B,C). The resulting deletions were relatively more precise than the *nd4*-targeted deletions, with 3/9 of the sequenced deletion junctions within both mito-nickase binding sites or intervening spacer sequence (Figure 2C,D).

**Figure 2.**
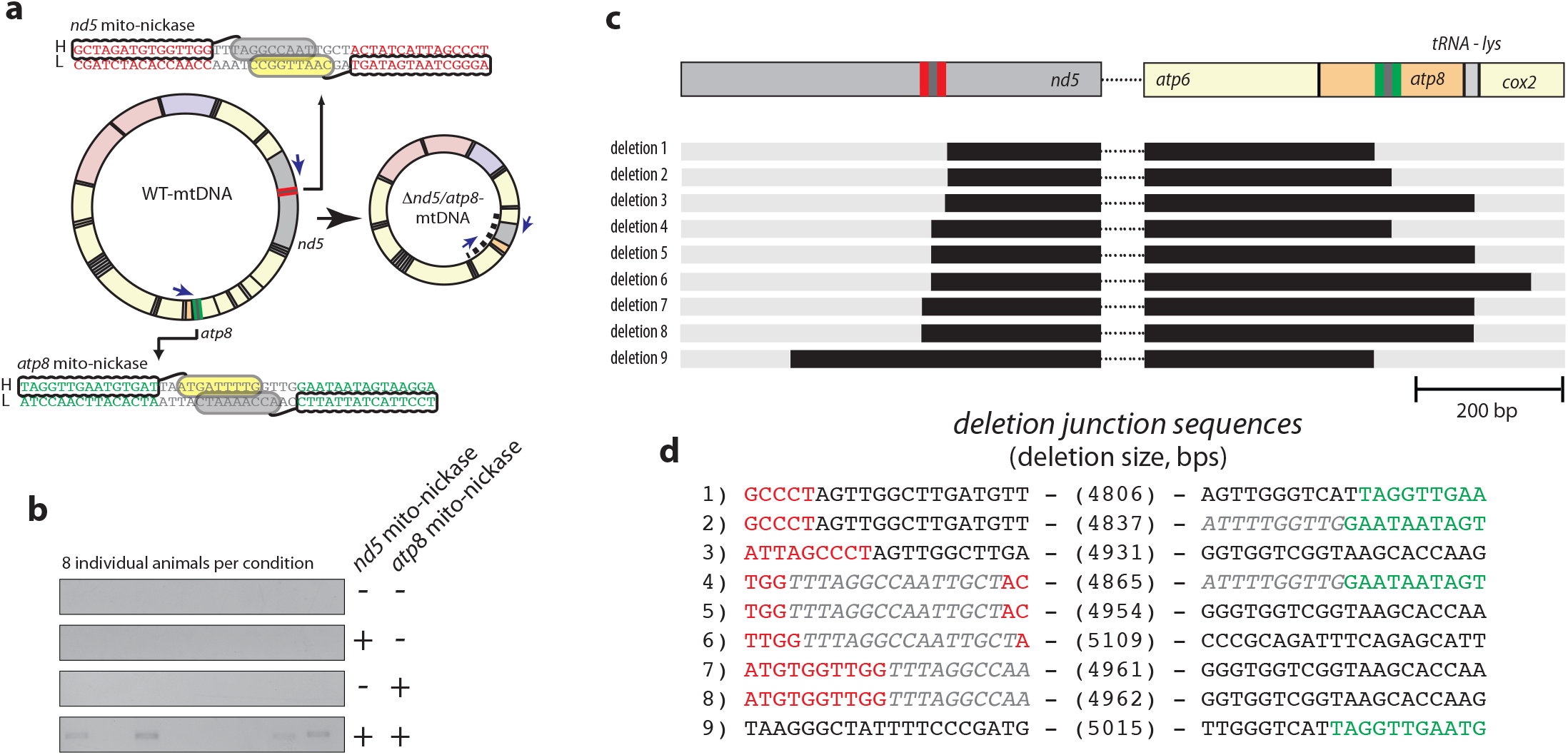
Multigenic deletion molecularly modeling the “common deletion” using mito-nickases. **a** Mito-nickases were targeted to *nd5* and *atp8*, with TALE binding and FokI activity on either the heavy (H) or light (L) strand. The active FokI is in yellow while the catalytically inactive variant, dFokI, is in grey. Deletions were detected using PCR with primers positioned outside of both nick sites (blue arrows). Expected PCR product indicated by dashed line. **b** Detection of deletions in animals injected with *nd5* and *atp8* mito-nickases. Every row contains 8 individual animals per condition. **c** Schematic of the regions around the *nd5* and *atp8* mito-nickase binding sites. Black bars indicate deletion regions. All mtDNA between *atp6* and *nd5* is deleted. Relative location of the *nd5* mito-nickase is in red, and *atp8* mito-nickase is in green. **c** Deletion junction sequences with the size of deletion in parentheses. In the left sequence panel, red sequences are the *nd5* mito-nickase binding sites and the grey, italicized sequence is the intervening spacer region. In the right sequence panel, green sequences are the *atp8* mito-nickase binding sites and the grey, italicized sequence is the intervening spacer region.

Nucleases are effective at shifting mtDNA heteroplasmy by selectively targeting and cutting genomes that contain the nuclease-binding site, thereby driving DNA degradation^8–12^. We asked whether this strategy could selectively target WT-mtDNA and preferentially enrich for the seeded Δ*nd5*/*atp8*-mtDNA genomes induced by nickases. A nuclease (mitoTALEN) was designed against *nd4*, located between the *nd5* and *atp8* mito-nickases target sites in WT-mtDNA (Figure 3A). The simultaneous delivery of the *nd5* mito-nickase, *atp8* mito-nickase, and the *nd4* mitoTALEN resulted in over 90% of injected animals harboring deletions (Figure 3B). These deletions were similar in sequence to nickase-injected animals (Figure 3E, S Figure 2). Using this approach, targeted deletions could also be generated in human 293T cells (Figure 3D, S Figure 3). Notably, we did not see the same variability in 293T mtDNA deletion products as we observed in zebrafish embryos (Figure 3D, S Figure 3). Zebrafish deletions were stable for months *in vivo*, as demonstrated by analyses of adult fish tissues (Figure 3C, S Figure 2). Interestingly, deletions were maintained in both eye and brain tissues, organs where the KSS deletion can manifest with phenotypic consequences for humans^26^ (Figure 3C, S Figure 4). Notably, we did not detect deletions in outcrossed embryos of female fish that maintained deletions in their fins, or in the ovaries of females that were positive for deletions in brain and eye tissue (Figure 3C). Interestingly, deletions were also observed after nuclease-alone treatment in zebrafish (Figure 3B) but not for 293T cells (Figure 3D). This could be due to the inefficiency of TALE transfection, as human cells require multi-plasmid transfection, but zebrafish can be directly microinjected. Alternatively, this could also be the result of differences between species-specific, double-strand break repair, or differences between a developing vertebrate embryo *in vivo* and human embryonic kidney cells *in vitro*.

**Figure 3.**
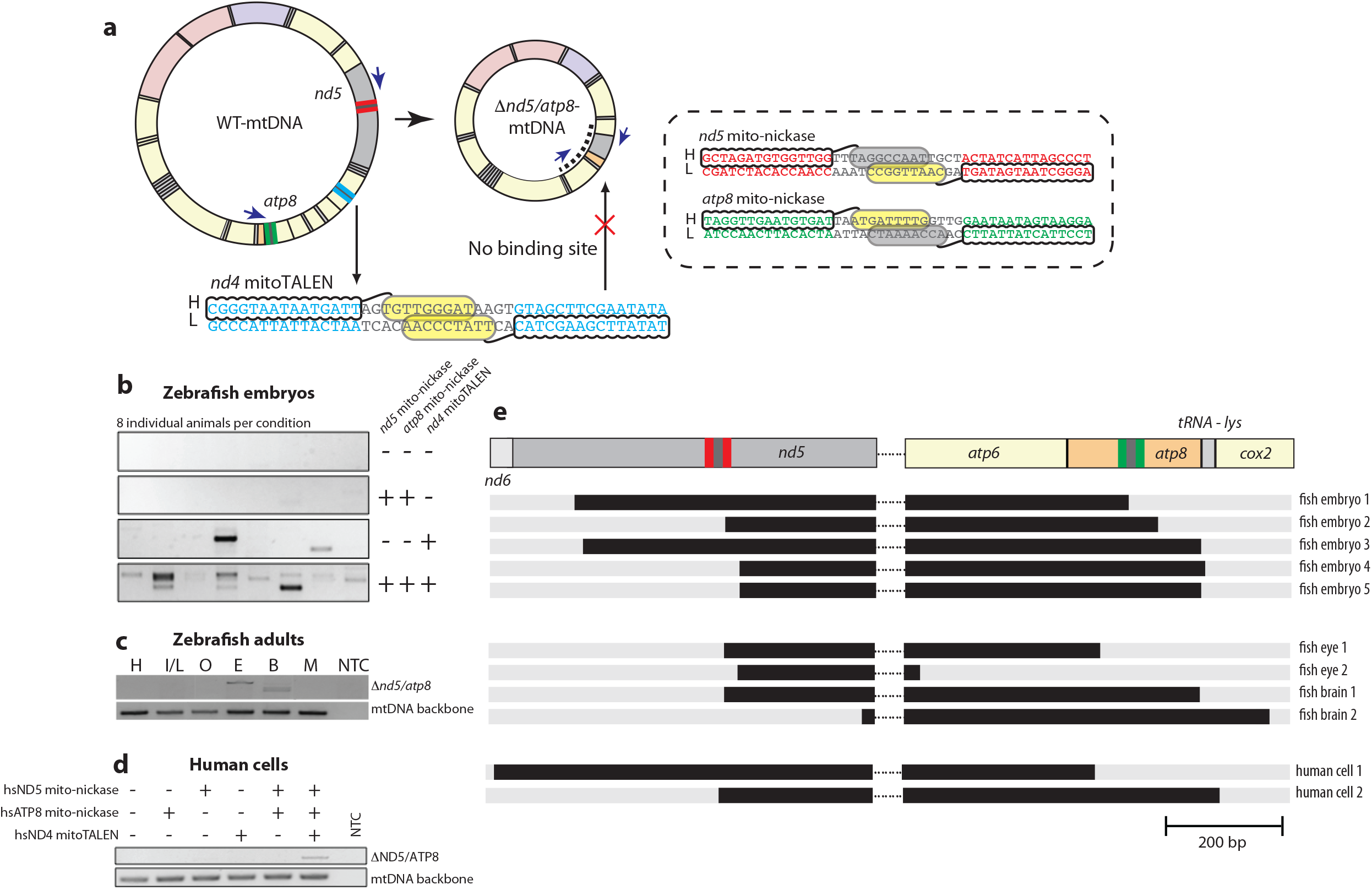
Expanding nickase-induced mtDNA deletions using a targeted mitoTALEN. **a** Schematic showing mitoTALEN targeting. A mitoTALEN targeting *nd4* was designed to selectively bind and cut the WT-mtDNA genome. The sequences targeted by the *nd5* and *atp8* mito-nickases are in the dotted box. **b** Detection of mtDNA deletions in injected zebrafish embryos. Every row contains 8 individual animals per condition. **c** Detection of mtDNA deletions in the organs of a four-month-old female zebrafish. H=heart, I/L=intestine/liver, O=ovaries, E=eyes, B=brain, M=Skeletal muscle. **d** Detection of mtDNA deletions in transfected human cells (293T cell line). **e** Schematic of the deleted regions around the *nd5* and *atp8* nickase binding sites. Black bars indicate deletion locations of mtDNA after editing by mito-nickases and mitoTALENs in zebrafish embryos, adult tissue, and human cells. All mtDNA between *atp6* and *nd5* is deleted. Relative location of the *nd5* mito-nickase is in red, and *atp8* mito-nickase is in green. NTC = no template control.

Deletions made using nickases did not usually fall within the predicted nick site and would at times be positioned just outside the TALE binding domains (Fig 1E, 2D, 3E). This observation, along with the strong enhancement of the mtDNA programming through the inclusion of mitoTALE nucleases (Figure 3B), suggested the potential for a novel molecular repair process. To test the role of nickases in mediating mtDNA programming, drop-out experiments were conducted by sequentially removing each component of the nickase. Surprisingly, each flanking mitoTALE-nickase could be simplified into a single mitoTALE arm (Figure 4A), so long as a nick was provided on the intervening mtDNA light strand (Figure 4B). The resulting deletions were reproducible, with single-nucleotide specificity noted (Figure 4C). Such reduced complexity of outcomes enabled the use of droplet digital PCR (ddPCR) to quantify mtDNA heteroplasmy across multiple embryos (Figure 4D). In a single seed step, a range of 1 in 2000 to 1 in 250 mtDNA molecules were altered (Figure 4D). To confirm that this effect applies to other regions of the mtDNA genome, a second deletion was generated between *coI* and *nd4* (Figure 4C), resulting in low complexity deletions with high precision at the nucleotide level (S Figure 5).

**Figure 4.**
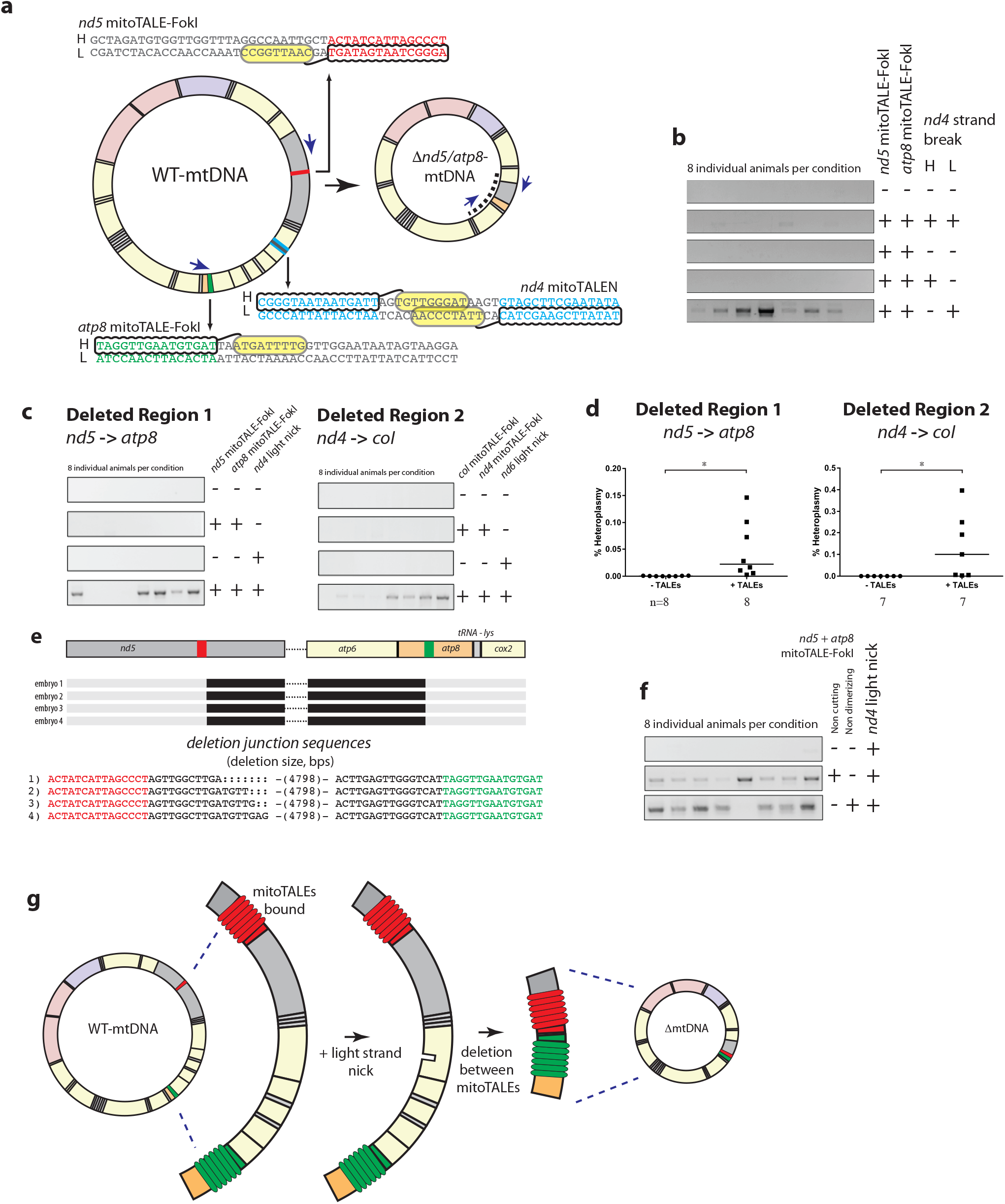
‘Block and nick’ strategy generates highly predictable mtDNA deletions. **a** Schematic showing mitoTALE arms (mitoTALE-FokI) positioned at *nd5* and *atp8* and a mitoTALEN positioned on *nd4*. **b** Detection of mtDNA deletions in zebrafish embryos injected with *nd5* and *atp8* mitoTALE-FokI with variable *nd4* heavy (H) and light (L) strand breaks **c** Detection of mtDNA deletions across two different multigenic regions, *atp8* to *nd5* and *nd4* to *coI*, using terminal mitoTALE-FokI binding and light strand nicking in-between. **d** Quantifying ΔmtDNA heteroplasmy of zebrafish embryos across two different multigenic regions. The median is shown by a horizontal bar. * = p < 0.05 **e** Schematic of the deleted regions around the *nd5* and *atp8* mitoTALE-FokI binding sites. Black bars indicate deletion locations of mtDNA deletions after mitoTALE-FokI and light strand nicking in zebrafish embryos. All mtDNA between *nd5* and *atp6* is deleted. Relative location of the *nd5* mitoTALE-FokI is in red, and *atp8* mitoTALE-FokI is in green. Deletion sizes are in parentheses, and additional single nucleotide deletions are shown with a colon (:). **f** Detection of mtDNA deletions in zebrafish after inhibiting either cutting (D450A amino acid substitution) or dimerization (p.D483A, p.R487A) of the terminal mitoTALE-FokI constructs. **g** Schematic of known molecular components involved in “block and nick” mtDNA editing. Every row in **b**, **c**, and **f** contains 8 individual animals per condition.

The naturally occurring 4977 bp deletion in human mtDNA has been recently reported to form from an unusual mechanism involving replication slippage around short sequence repeats that flank the deletion breakpoint^27^. The results noted here suggest an alternative mechanism; no apparent sequence repeats flanking *de novo* mtDNA deletions are noted in these induced mtDNA deletions that would be indicative of replication slippage or microhomology-mediated end joining break repair^28^. Nicking the mtDNA heavy strand adjacent to one of the repeat sequences reportedly enriches the ‘common deletion’ observed in humans. However, the inverse was noted here: by nicking the mtDNA light strand, we could generate designer mtDNA variants (Figure 4B). Additionally, we found no loss of activity when we eliminated the cutting or dimerization ability of the terminal TALE-FokI proteins(Figure 4F). This, combined with the sequence data confirming that deletions are made adjacent to the TALE target sequence (Figure 4E, S Figure 5), suggests these targeted mtDNA binding proteins are acting as a blockade to mitochondrial mtDNA machinery that is triggered by nicking the light strand. Although the precise nature of the underlying mechanism of induced mtDNA deletions is still to be elaborated, the requirement for terminal targeting proteins surrounding an intervening light strand nick suggests a ‘block and nick’ working model for mtDNA targeted deletions (Figure 4G).

The ability to precisely manipulate the mitochondrial genome is critical to our understanding of human disease and the biology of this complex organelle. Mitochondrial gene editing can be approached as a population genetics problem as thousands of mtDNA genomes are found within each cell. Hence, the ‘block and nick’ mutation process described herein can address the first step of mtDNA genome manipulation by ‘seeding’ mtDNA sequences that harbor desired variants (Figure 5). The most apparent limitation of this technology is the low initial heteroplasmy levels observed following this initial seed step. By elucidating the mechanism used to generate these low-level, high-precision deletions, we hope to increase the yield of mtDNA-edited products. The subsequent ability to use specific nucleases to drive heteroplasmy levels based on unique nucleotide sequence also provides one potential approach to expand mtDNA deletions to biologically significant heteroplasmy levels (Figure 5). Such an approach would enable the tunable production of cells with differential heteroplasmy levels to recapitulate phenotypes found in human disease, a critical phenomenon known as the threshold effect^29–31^. Similarly, the development of complex *in vivo* animal models that mirror human disease (from zebrafish to mammals) would be possible through the temporal or spatial (i.e. organ-specific) regulation of mitoTALENs or other targeted nucleases after a one-time delivery of “block and nick” reagents to seed targeted mtDNA deletions in the embryo, as conducted here in the zebrafish test system. To our knowledge, the system described here is the first user-guided system to address direct sequence modification of mtDNA *de novo*. We believe these findings can accelerate mitochondria research and lead to insights and improved therapies for mtDNA disease.

**Figure 5.**
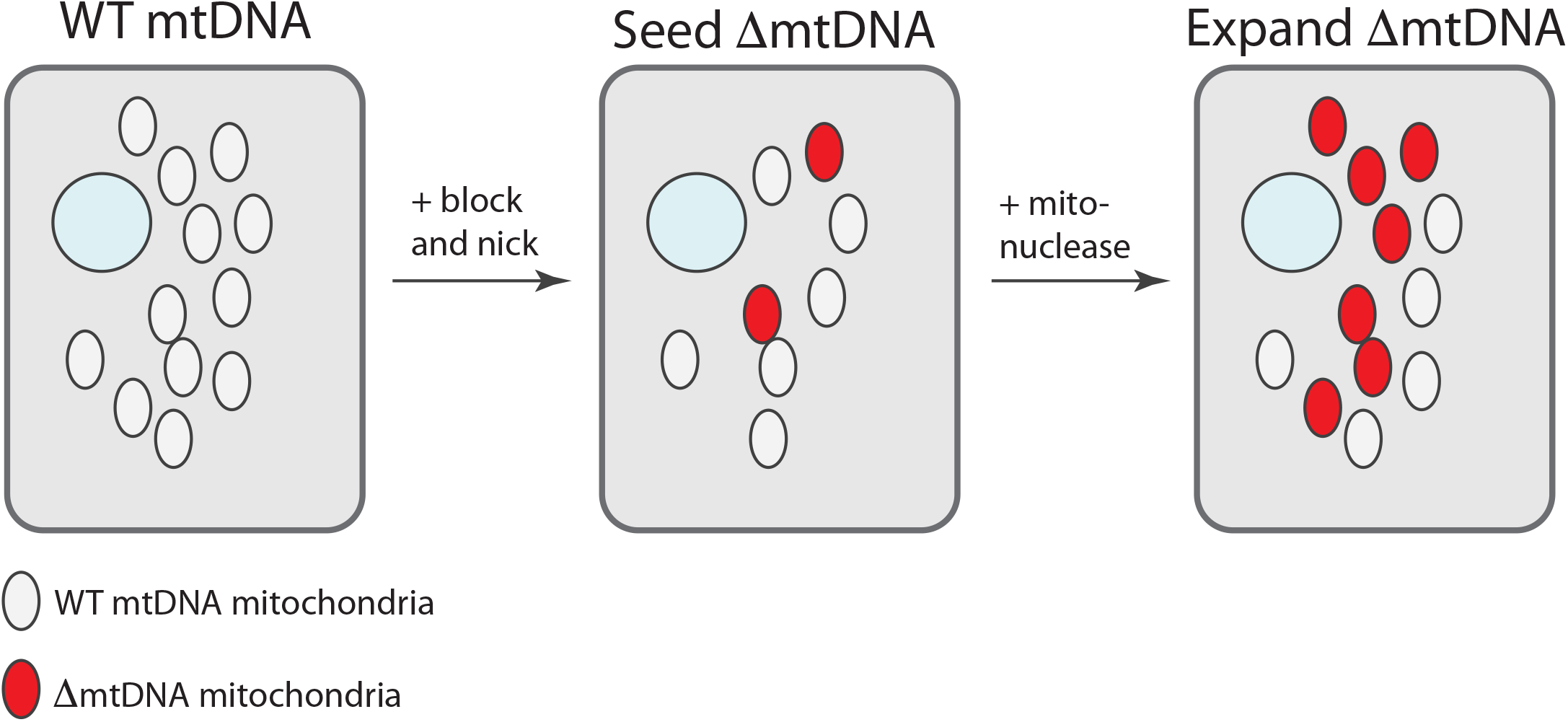
Cell- and tissue-level mtDNA population engineering toolkit using “seed and expand”. Targeted deletions are seeded in the mitochondrial network using targeted block and nick technology. These designer mitochondrial genomes are separately expanded through heteroplasmic shift from targeted mito-nucleases.

## Materials and methods

### Zebrafish handling

All animal work was conducted under Mayo Clinic’s institutional animal welfare approval.

### MitoTALE-nickase and -nuclease assembly

The nuclear localization sequence of GoldyTALEN was replaced in the expression plasmids pT3TS-GoldyTALEN (pT3TS-mitoTALEN, T3 promoter mRNA expression plasmid^32^) and pC-GoldyTALEN (pC-mitoTALEN miniCaggs promoter expression plasmid) by the mitochondrial targeting sequence of zebrafish *idh2* (S Figure 1). To generate a mitoTALE-nickase, the p.D450A amino acid substitution was made in the FokI domain of one half of a TALEN pair^23^. For a non-cutting TALEN, two TALENs harboring the p.D450A substitution were used. To make a non-dimerizing FokI, two amino acid substitutions were made (p.D483A, p.R487A^33^) and each TALE arm harbored these variants in *cis*. The final mitoTALE backbone plasmids used in this paper will be made available at Addgene. To target a specific sequences in the genome, RVDs were cloned into the GoldyTALE backbone using the FusX assembly system^18^. The RVDs used were HD=C, NN=G, NI=A, NG=T. The sequences targeted by all TALEs in this manuscript can be found on S Table 1.

### Zebrafish TALE mRNA embryo injections and deletion screening

Once the RVDs were cloned, mRNA was made for zebrafish microinjections. Each TALEN expression plasmid was digested with SacI for 2-3 hours at 37°C, then synthetic mRNA was made (T3 mMessage mMachine kit, Ambion) and extracted using a phenol/chloroform extraction (according to T3 mMessage mMachine kit instructions). 12.5 pg of each mitoTALE-nickase (Figures 1–3) or mitoTALE-FokI arm (Figure 4) was co-injected into one-cell embryos, with 50 pg of mitoTALEN (Figures 1–3) or mitoTALE-nickase (Figure 4) when indicated. After 3 days, individual larval zebrafish were collected in 1.7 mL microcentrifuge tubes, and DNA was extracted in 40 uL of an extraction buffer (50 mM Tris– HCl pH 8.5, 1 mM EDTA, 0.5% Tween-20, 200 μg/ml proteinase K) at 55°C overnight and centrifuged at 17,000xg for 1 minute after extraction was complete. In each condition (control and experimental), 8 individual zebrafish embryos were collected and screened for mtDNA deletions. Replicates were done in triplicate for all mtDNA editing experiments in both zebrafish embryos and human cells with the exception of Figure 1C, Figure 4B and Figure 4F which were done in duplicate. One microliter of DNA extract was then used for PCR (Platinum Taq, Invitrogen). All PCR reactions were kept on ice until the thermocycler reached 95°C. Primer concentrations were 0.4 uM for all reactions, and the final reaction volume was 25 uL. To include MgCl_2_ and dNTPs, and loading dye, the 5x MyTaq Red buffer was used (Bioline Cat# BIO-37112). For screening *nd4* deletions, the following thermocycler parameters were used: 95°C for 5’, (95°C for 30” - 60°C for 30” - 72°C for 1’) x 35, 72°C for 5’. For screening zebrafish *nd5/atp8* deletions, the following thermocycler parameters were used: 95°C for 5’, (95°C for 30” - 64°C for 30” - 72°C for 1’) x 35, 72°C for 5’. For control PCR to detect successful DNA extraction, the following thermocycler parameters were used: 95°C for 5’, (95°C for 30” - 55°C for 30” - 72°C for 1’) x 35, 72°C for 5’. In all cases 5 uL of PCR product was run on a 1-1.5% gel and extracted for sequencing using the Qiaex II gel extraction kit (Qiagen), then cloned TOPO TA cloned into a sequencing vector using the TOPO PCR cloning kit (Invitrogen Cat# K4575J10) and Sanger sequenced.

### Human mitoTALE transfections and deletion screening

293T cells (ATCC CRL-3216) were transfected with 500 ng of each pC-mitoTALE-nickase and 1 ug of pC-mitoTALEN when indicated. 500 ng of a plasmid containing EGFP was included to ensure proper transfections and to act as a negative control. Transfections were done using the 100uL tips of the Neon transfection system (Invitrogen) and electroporation conditions were as recommended by Invitrogen for HEK293 cells (Pulse voltage = 1,100 v, pulse width = 20 ms, pulse number = 2, cell density ~ 5 x 10). Cells were immediately put into 6-well plates post transfection in DMEM media containing 10% FBS, pen-strep, 50 ug/mL uridine, and 1mM sodium pyruvate. After 2 days, cells were moved to a T75 flask to allow for expansion. After 6 days cells were collected and mtDNA extraction was performed using a mitochondrial DNA isolation kit (Biovision Cat# K280-50). One modification was made to the protocol; on step 6 of the Mitochondrial DNA Isolation Protocol from the user manual, we centrifuged at 2,000xg for 6 minutes at 4°C instead of 700xg for 10 minutes at 4°C. The resulting DNA was diluted to 50 ng/uL and 1uL of this was used for PCR to detect mtDNA deletions (with the same reagents as zebrafish PCR). Primer sequences used can be found on S Table 2 and concentrations were 0.4 uM per reaction, with a final reaction volume of 25 uL. Importantly, taq polymerase was not added to the reaction until the reaction reached 95°C. Thermocycler parameters to detect deletions were as follows: 95°C for 5’, (95°C for 30” - 64°C for 30” - 72°C for 1’) x 35, 72°C for 5’. For control PCR to detect successful DNA extraction, the following thermocycler parameters were used: 95°C for 5’, (95°C for 30” - 55°C for 30” - 72°C for 1’) x 35, 72°C for 5’. After amplification 5 uL of PCR product was run on a 1.5% gel and extracted for sequencing using the Qiaex II gel extraction kit (Qiagen Cat#20051), then cloned TOPO TA cloned into a sequencing vector using the TOPO PCR cloning kit (Invitrogen Cat# K4575J10) and Sanger sequenced.

### Zebrafish organ dissection and DNA extraction

Organ dissection was conducted on six euthanized 4-month-old zebrafish to collect the heart, liver, intestine, eye, brain, skeletal muscle and ovaries^34^. Cont’rols (animals not injected with mitoTALEs) were age- and sex-matched, and their organs collected at the same time. In total, two control fish were dissected (one male and one female) and four fish injected with mitoTALEs were dissected (two males, and two females). Organs were added to 100 uL of a DNA extraction buffer (50 mM Tris– HCl pH 8.5, 1 mM EDTA, 0.5% Tween-20, 200 μg/ml proteinase K) and incubated at 55°C overnight, followed by centrifugation at 17,000xg for 1 minute. Bridge PCR as well as control PCR to ensure proper DNA extraction was performed using 2 uLs of extract in a 25 uL reaction, with conditions otherwise identical to the embryo screening of *nd5/atp8* deletions. Sanger sequencing was conducted following PCR to identify deletion junctions

### Droplet Digital PCR (ddPCR) to measure heteroplasmy

Droplet digital PCR was performed on DNA extracted from both zebrafish embryos and adult tissue. To detect heteroplasmy, one amplicon and TaqMan probe containing a 5’HEX fluorophore and 3’quencher (IDT) was designed against a common region in the mtDNA genome (*nd1*) and another amplicon and probe containing a 5’FAM fluorophore and 3’ quencher were designed to bridge deletion (ΔmtDNA) breakpoint. To ensure proper droplet saturation for a quantitative measurement of wild type genomes using the HEX fluorophore, a dilution of 1:100 was performed and accounted for upon data analysis. To detect the deleted mtDNA genomes using the FAM fluorophore, no dilution was performed. 1uL of sample was added to 10uL of the 2x ddPCR supermix for probes (BioRad Cat#186-3023) along with 900nM of PCR primers and 250nM of probe, 1uL of HindIII, and water to a final volume of 20uL. Samples were loaded into a 96-well plate and heat-sealed with tinfoil, vortexed, and briefly centrifuged. Droplets were generated on a BioRad AutoDG automatic droplet generator. After droplet formation, samples were moved to a thermocycler and PCR was performed with 95°C for 10’, (94°C for 30” - 60°C for 1’) x 40, 98°C for 10’, and 4°C hold until the samples were read on a QX200 droplet reader. Droplet analysis was performed with QuantaSoft software. The copy number of mtDNA molecules in 1uL of sample for both wild type and deleted genomes were determined to calculate heteroplasmy. For this, the following equation was used: ([ΔmtDNA]/[*nd1*]) x 100, where brackets are the concentration. Droplet digital PCR was performed with technical duplicates and averaged to reach each individual data points.

Either 7 or 8 animals were screened for heteroplasmy quantification in zebrafish embryos (Figure 4D), or 2 for adult tissue (S Figure 4B). Animals were chosen based on a positive signal for bridge PCR along with the same number of uninjected control animals. Once heteroplasmy was determined, GraphPad PRISM software was used to create dot plots and identify either the median (Figure 4B) or the mean (S Figure 4B) and to calculate statistical significance (P value) using a two-sided Mann-Whitney U test for Figure 4B.

## Acknowledgements

This work was supported by a gift from the Marriott Foundation and NIH grants GM63904, HG 006431, and P30DK084567 to SCE. We thank Drs. Eric Schon, Vamsi Mootha, Michio Hirano, and José Antonio Enríquez for advice. We also thank the team at B-MoGen Biotechnologies for advice on mtDNA extraction protocols. Finally, we thank Dr. Katherine Campbell, Roberto Lopez Cervera, and Zachary WareJoncas for critical review of this manuscript.

## Author contributions

The paper was written by JMC and SCE and edited by EPC and KJC. Experiments were executed by JMC, HA and NVF with experimental guidance from SCE, TJN, KJC, DO and EPC. Data analysis was completed by JMC, EPC, NVF and SCE. WL and XX identified the zebrafish *idh2* locus and the putative mitochondrial targeting sequence from the protein trap gene-breaking transposon lines.

## Financial disclosures

Mayo Clinic, JMC and SCE have a financial interest related to this research through a license to B-MoGen Technologies, Inc.

