## Supplementary Materials for "Engineering targeted deletions in the mitochondrial genome"

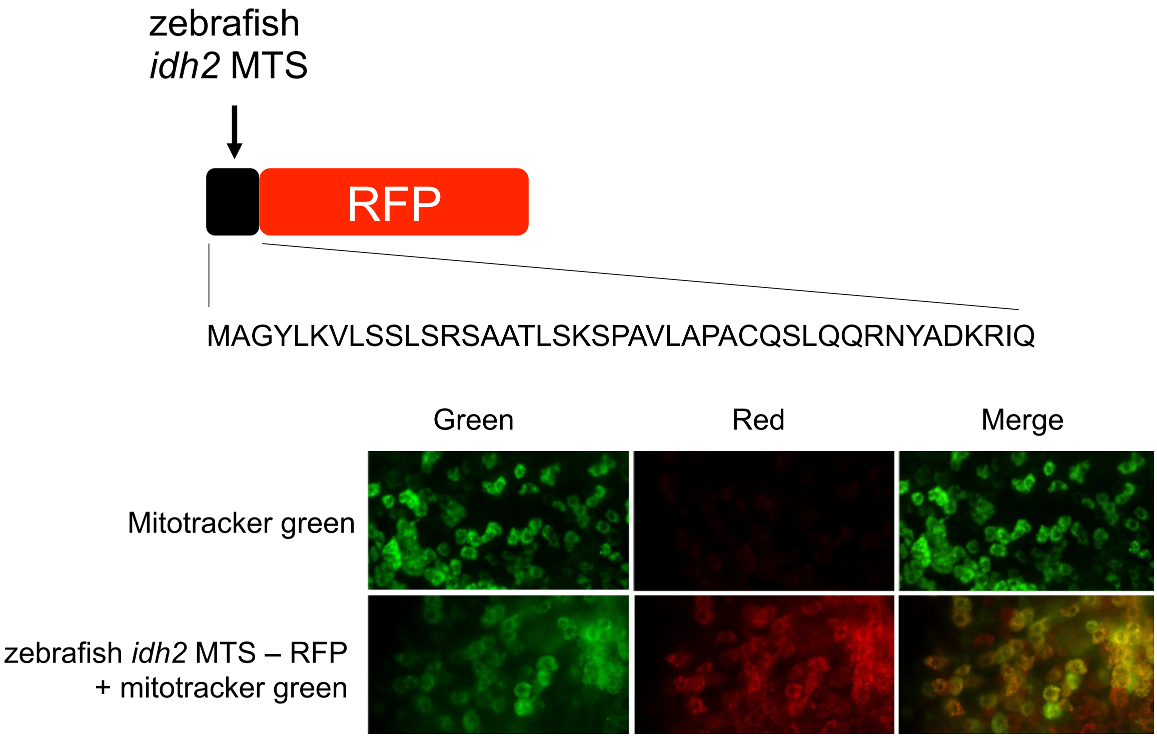


**Supplemental Figure 1 |** Characterization of the zebrafish isocitrate dehydrogenase 2 (*idh2*) mitochondrial targeting sequence (MTS). The MTS from the transgenic protein-trap zebrafish line GBT268 was characterized using MitoProt and fused to RFP for characterization in zebrafish embryos. One-cell embryos were injected with mitotracker green, both with and without the *idh2*MTS-RFP fusion. After 9 hours embryos were imaged on a Zeiss Lightsheet Z.1 single plane illumination microscopy (SPIM) microscope.

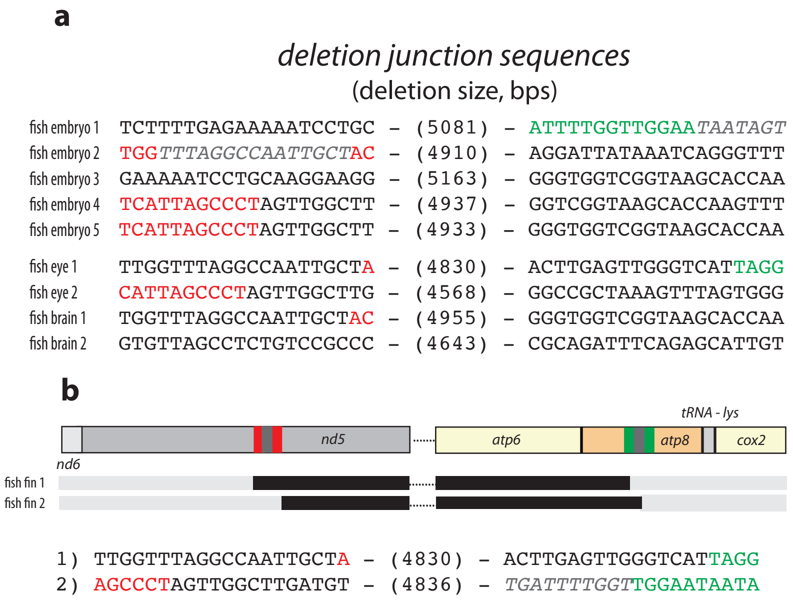


**Supplemental Figure 2 |** Deletion junction sequences in zebrafish injected with the *nd5/atp8* mito-nickases and *nd4* mitoTALEN outlined in Figure 3A. In the left sequence panel, red sequences are the *nd5* nickase binding sites and the grey, italicized sequence is the intervening spacer. In the right sequence panel, green sequences are the *atp8* nickase binding sites and the grey, italicized sequence is the intervening spacer. **a** Sequences of deletion junctions corresponding to Figure 3E. **b** mtDNA deletions from fin biopsies of two four-month-old zebrafish (one male, one female). Schematic of the deleted regions around the *nd5* and *atp8* nickase binding sites. Black bars indicate deletion locations of mtDNA after editing by mitoTALE-nickases and mitoTALENs in zebrafish embryos, adult tissue, and human cells. All mtDNA between *atp6* and *nd5* is deleted. Relative location of the *nd5* nickase is in red, and *atp8* nickase is in green.

**a**

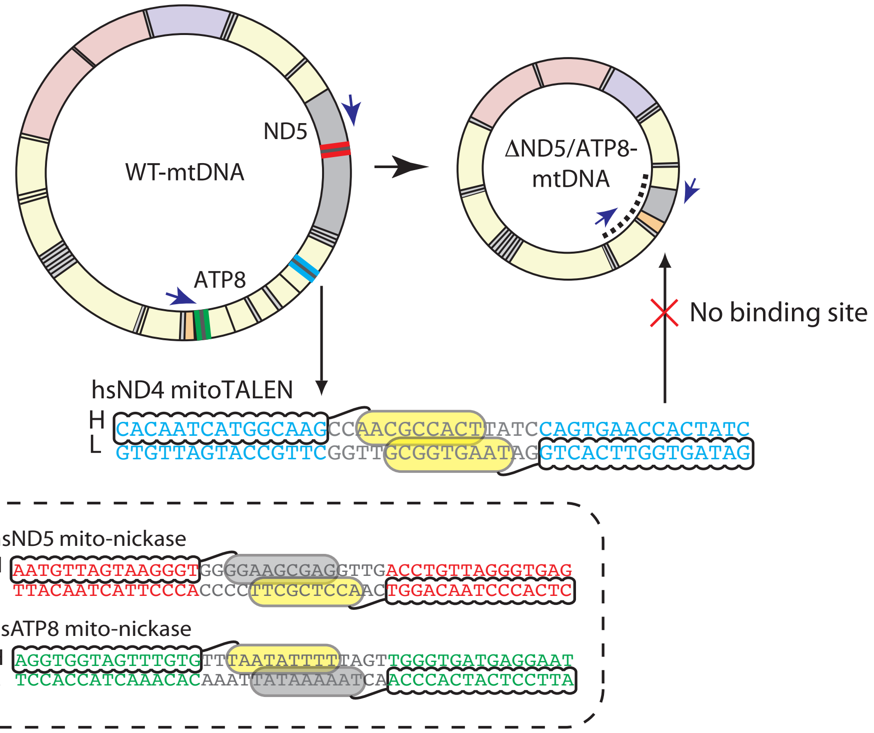


**b**

*deletion junction sequences*
(deletion size, bps)

1) GGT*GGGGAAGCGAGGTTG*AC – (4806) – CGGGCCCTATTTCAAAGATT
2) TAGTAGTAGTTACTGGTTGA – (4826) – ATAATTTTTTATTTTTTATGG

**Supplemental Figure 3 |** Targeted mtDNA deletions in human cells to molecularly model the common deletion. **a** Schematic showing mito-nickase and mitoTALEN targeting. A mitoTALEN targeting ND4 was designed to selectively bind and cut the WT-mtDNA genome. The sequences targeted for the ND5 and ATP8 nickases are in the dotted box. **b** Deletion junction sequences from schematic in Figure 3E. In the left sequence panel, red sequences are the *nd5* nickase binding sites and the grey, italicized sequence is the intervening spacer.

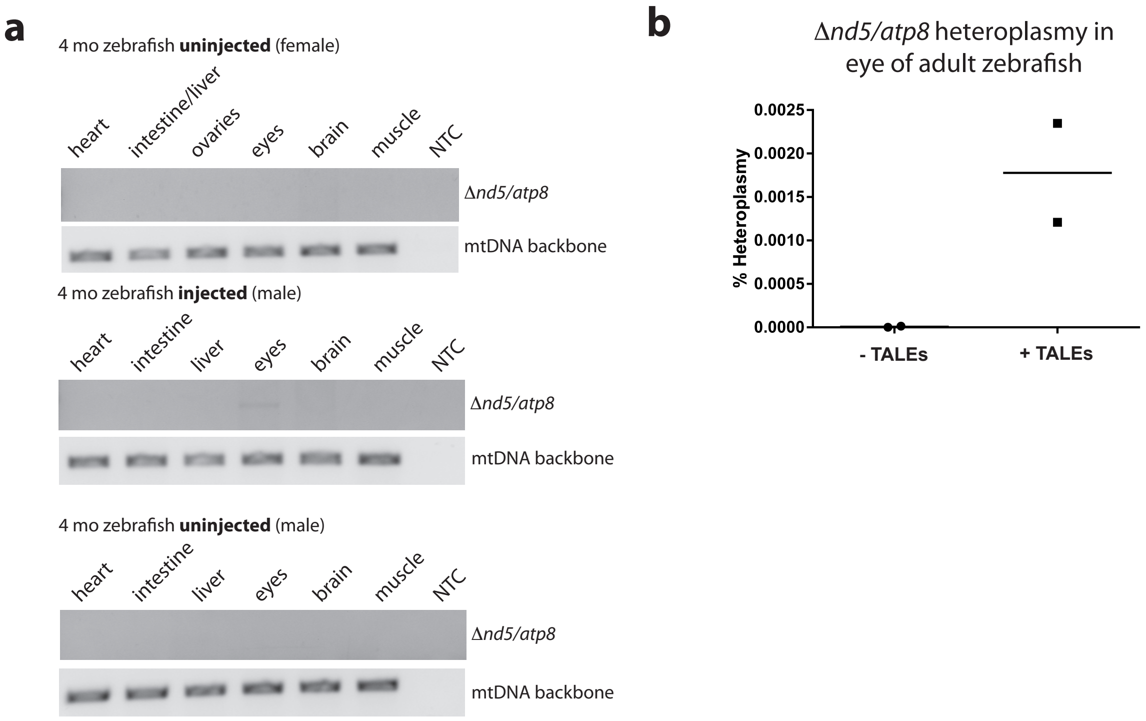


**Supplemental Figure 4 | Targeted mtDNA deletions in four-month-old zebrafish. a** Detection of *nd5* to *atp8* mtDNA deletions in various organs of zebrafish injected with the *nd5* and *atp8* mito-nickases and the *nd4* mito-TALEN that can be found in Figure 3A. NTC=no template control. **b** Heteroplasmy levels in the eyes of zebrafish injected with and without the mito-nickases and mitoTALEN (one male and one female from each group). The mean is shown by a horizontal bar.

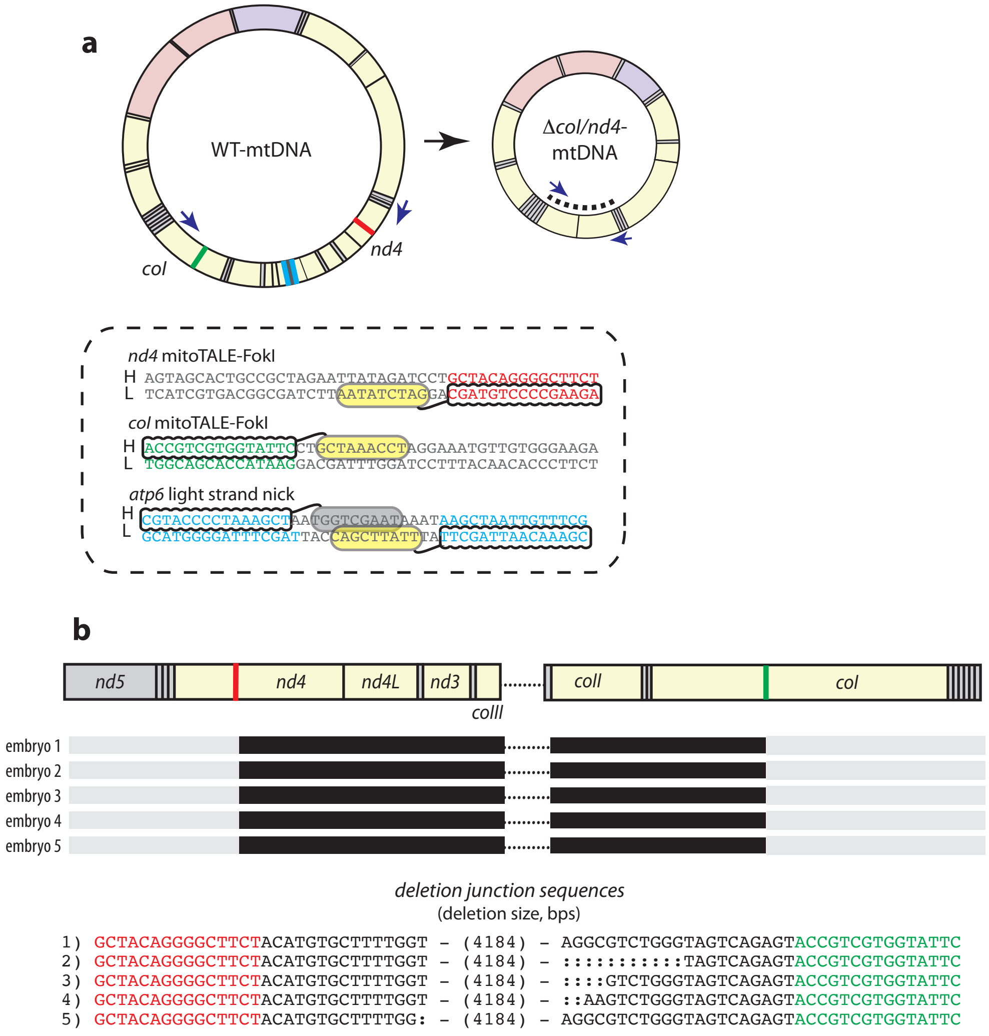


**Supplemental Figure 5 | ** Zebrafish *nd4* to *co1* deletions (Figure 4, Region 2) made using the 'block and nick' strategy **a** Schematic showing single TALE arms (mitoTALE-FokI) positioned at *nd4* and *co1* and a mitoTALE-nickase cutting the light strand positioned on *nd6*. TALE binding sequences are in the dotted box. **b** Schematic of the deleted regions around the *nd4* and *coI* mitoTALE-FokI binding sites. Black bars indicate deletion locations of mtDNA deletions after mitoTALE-FokI and light strand injections in zebrafish embryos. All mtDNA between *coII* and *coIII* is deleted. Relative location of the *nd4* mitoTALE-FokI is in red, and *co1* mitoTALE-FokI is in green. Deletion sizes are in parentheses, and additional single nucleotide deletions are shown with a colon (:).

| | S Table 1: TALE mtDNA binding targets | | |
|---|---|---|---|
| | | Left (5’-3’) | Right (5’-3’) |
| zebrafish | *nd4* mito-nickase (1) | AAGTTCTGGTTTTAA | TTACACCTAATTCCT |
| | *nd4* mito-nickase (2) | ACGAGTTAGAAGCAT | GACTAAAATGAACCT |
| | *nd5* mito-nickase | GCTAGATGTGGTTGG | AGGGCTAATGATAGT |
| | *atp8* mito-nickase | TAGGTTGAATGTGAT | TCCTTACTATTATTC |
| | *nd4* mitoTALEN | CGGGTAATAATGATT | TATATTCGAAGCTAC |
| | *nd4* TALE-FokI | AGAAGCCCCTGTAGC | |
| | *coI* TALE-FokI | ACCGTCGTGGTATTC | |
| | *atp6* mito-nickase | CGTACCCCTAAAGCT | CGAAACAATTAGCTT |
| human | hsND5 mito-nickase | AATGTTAGTAAGGGT | CTCACCCTAACAGGT |
| | hsATP8 mito-nickase | AGGTGGTAGTTTGTG | ATTCCTCATCACCCA |
| | hsND4 mitoTALEN | CACAATCATGGCAAG | GATAGTGGTTCACTG |

| | S Table 2: Primers and Probes | | |
|---|---|---|---|
| | | Forward (5’-3’) | Reverse (5’-3’) |
| zebrafish | *nd4* deletion<br>*bridge PCR primers* | CCAGGTGATGAATAAGGCGATTGAGGT | CCAGGTGATGAATAAGGCGATTGAGGT |
| | *nd5* -> *atp8* deletion<br>*bridge PCR primers* | GCAATAATGCTTCCTCAGGCAAGCCGT | GACGCGGTACCCGGACGATTAAACCA |
| | *nd4* -> *coI* deletion<br>*bridge PCR primers* | CGGATGAGGTTAGTCCGTGGGCGA | ACCCGAGCATACTTCACATCCGCC |
| | mtDN*A control primers*<br>(zebrafish MT-TL1) | TGATTGTAACAGTCCTCGGGGGCC | AGGATCGGAAAAAGGGGGCCCATAC |
| | *nd5* -> *atp8* deletion<br>*ddPCR primers* | TGCTAGATGTGGTTGGTTTAGGC | CATTCAACCTAATGACCCAACTCAAG |
| | *nd5* -> *atp8* deletion<br>*ddPCR probe (FAM)* | TTGCTACTATCATTAGCCCTAGTTGGCTTG | |
| | *nd4* -> *coI* deletion<br>*ddPCR primers* | GCACTGCCGCTAGAATTATAGATCC | TCTCACATTCTTCCCACAACATTTCC |
| | *nd4* -> *coI* deletion<br>*ddPCR probe (FAM)* | TACCGTCGTGGTATTCCTGCTAAACC | |
| | *nd5* -> *atp8* deletion<br>*ddPCR (eye) primers* | ATTCAACCTAATGACCCAACTCAAG | GCCCCTGAGCATAAGAATAATATGG |
| | *nd5* -> *atp8* deletion<br>*ddPCR (eye) primers*<br>*(FAM)* | TCCACATCTGCACTCACGCTT | |
| | *nd1* ddPCR<br>*primers* | CCCCTCTGTAAGATCGAACGG | ACTTCCACTATGACCATTAGCCA |
| | *nd1* ddPCR<br>*probe (HEX)* | TCGGTTTGTTTCTGCCAGCGTTG | |
| human | *nd5* -> *atp8* deletion<br>*bridge PCR primers* | GCGATGGAGGTAGGATTGGTGCTGTGG | CTCATGAGCTGTCCCCACATTAGGCTT |
| | mtDNA *control primers*<br>*(*human MT-TL1) | GAGTTTTATGGCGTCAGCGA | GGACAAGAGAAATAAGGCCTACTTC |
